# Escaping inbreeding: the demographic path to genetic recovery

**DOI:** 10.1101/2025.05.20.655063

**Authors:** Collin W. Ahrens, Adam D. Miller, Luke W. Silver, Elspeth A. McLennan, Carolyn J. Hogg, Andrew R. Weeks

**Affiliations:** Cesar Australia, Brunswick, Victoria, Australia; Western Sydney University, Richmond, NSW, Australia; College of Sciences, Flinders University, Bedford Park, South Australia; Australasian Wildlife Genomics Group, The University of Sydney, Sydney, Australia; School of BioSciences, The University of Melbourne, Parkville, Australia

## Abstract

Bottlenecks pose a major threat to species persistence by reducing genetic diversity and increasing inbreeding. Although evolutionary theory suggests these constraints can be overcome, empirical evidence has largely come from invasive species. Here, we analyse whole-genome data from 418 koalas (*Phascolarctos cinereus*) across 27 populations to reconstruct demographic histories and examine rare genetic variation. We find that northern populations retain higher genetic diversity, while southern populations, despite severe bottlenecks, exhibit larger and increasing effective population sizes (*N*_e_). This apparent contradiction is explained by increased reshuffling of genetic variation through recombination, and the accumulation of rare alleles during recent demographic expansion. Our findings demonstrate that rapid population growth can substantially elevate *N*_e_, offering an evolutionary pathway by which threatened populations may “escape” the genetic risks of inbreeding.

## Introduction

Populations that undergo prolonged bottlenecks experience a cascade of genetic consequences (*1*). Severe reductions in population size increase the effects of random genetic drift, leading to the loss of genetic diversity and the fixation of deleterious alleles (*2*). Inbreeding becomes inevitable, elevating the risk of inbreeding depression as segregating recessive deleterious mutations are expressed at higher frequencies (*3*). This accumulation of ‘genetic load’ can drive negative changes in fitness-related traits, including reduced fertility (*4*) and survival rates (*5*), along with increased susceptibility to disease (*6*). Beyond direct fitness consequences, the erosion of genetic variation limits a population’s adaptive potential, constraining its ability to respond to environmental change (*7,8*). As a result, bottlenecked populations often face an extinction vortex, where reduced genetic diversity and declining population sizes reinforce one another in a feedback loop of demographic and evolutionary decline (*9*).

While prolonged bottlenecks are often associated with irreversible genetic loss (*10,11*), rapid demographic expansion can generate increased recombination and facilitate the accumulation of novel mutations (*12*). This can reduce the immediate effects of genetic drift by increasing effective population size (*N*_e_), thereby slowing diversity loss and mitigating the risk of inbreeding depression (*13*). A striking parallel to this dynamic is seen in the *genetic paradox of invasive species*, which often thrive despite undergoing severe founder events started from a small number of individuals (*14*). Upon introduction to a new environment, invasive populations frequently pass through genetic bottlenecks but subsequently undergo rapid population growth, allowing them to escape the expected consequences of genetic drift and inbreeding (*14*). This paradox suggests demographic expansion is a key mechanism for recovering genetic diversity. Whether this process can also facilitate recovery in threatened species remains unclear, but it challenges the current paradigm that assumes low genetic diversity constrains their persistence and evolutionary potential.

The koala (*Phascolarctos cinereus*) is an iconic arboreal marsupial uniquely adapted to a eucalypt-based diet. Once widespread across eastern and southern Australia, koalas have experienced dramatic population declines driven by habitat loss, disease, and historical hunting pressure (*15*). Northern populations in Queensland (QLD), New South Wales (NSW), and the Australian Capital Territory are now listed as Endangered under the Australian Federal Environment Protection and Biodiversity Conservation (EPBC) Act 1999. In contrast, southern populations in Victoria (VIC) remain abundant and subject to intensive management (*15*). This management traces back to one of the longest-running conservation translocation programs in history, initiated in the 1890s when a small number of koalas were relocated to French Island (VIC) to safeguard against losses from the fur trade (*16*). Throughout the 20^th^ century, individuals from this refuge were reintroduced to mainland Victoria, facilitating a remarkable demographic recovery (*15*). Koala numbers in Victoria subsequently surged to more than half a million individuals (*17*), leading to issues of overabundance and necessitating management interventions such as translocation, fertility control, and, at times, culling (*15*). Given this unique history, southern koalas present a compelling case study for understanding how rapid demographic expansion may mitigate the effects of severe bottlenecks and inbreeding.

As part of koala conservation efforts, whole-genome sequencing (WGS) was performed on 418 individuals from across their range (*18*), representing one of the largest WGS datasets for a threatened species. Here, we make use of this novel dataset to reconstruct the demographic histories of koala populations and to determine how they have shaped the contemporary distribution of genetic variation. Specifically, we explore how population bottlenecks and subsequent demographic expansion have influenced effective population size (*N*_e_), levels of inbreeding, recombination and the accumulation of low-frequency alleles—key factors that may determine whether a species can escape the genetic constraints of a bottleneck.

### Demographic history of koala populations

To explore the demographic history of koala populations, we employed population-level coalescent models (SMC++; *19*) . These analyses revealed a significant population expansion approximately 10,000 generations ago, broadly consistent with earlier findings (*20*). However, notable differences emerged thereafter among state meta-populations. The VIC meta-population showed a continued decline around 1,000 generations ago (Fig. 1H), contrasting with the QLD and NSW meta-populations, which stabilized above an *N*_e_ of 1,000 (Fig. 1H). These data suggest that a substantial decline in the VIC meta-population predates European colonization (circa 16-1700s; Figs. 1B,E) and that European hunting for the fur trade was not the only cause of their near extirpation in Victoria (*15*).

**Figure 1.**
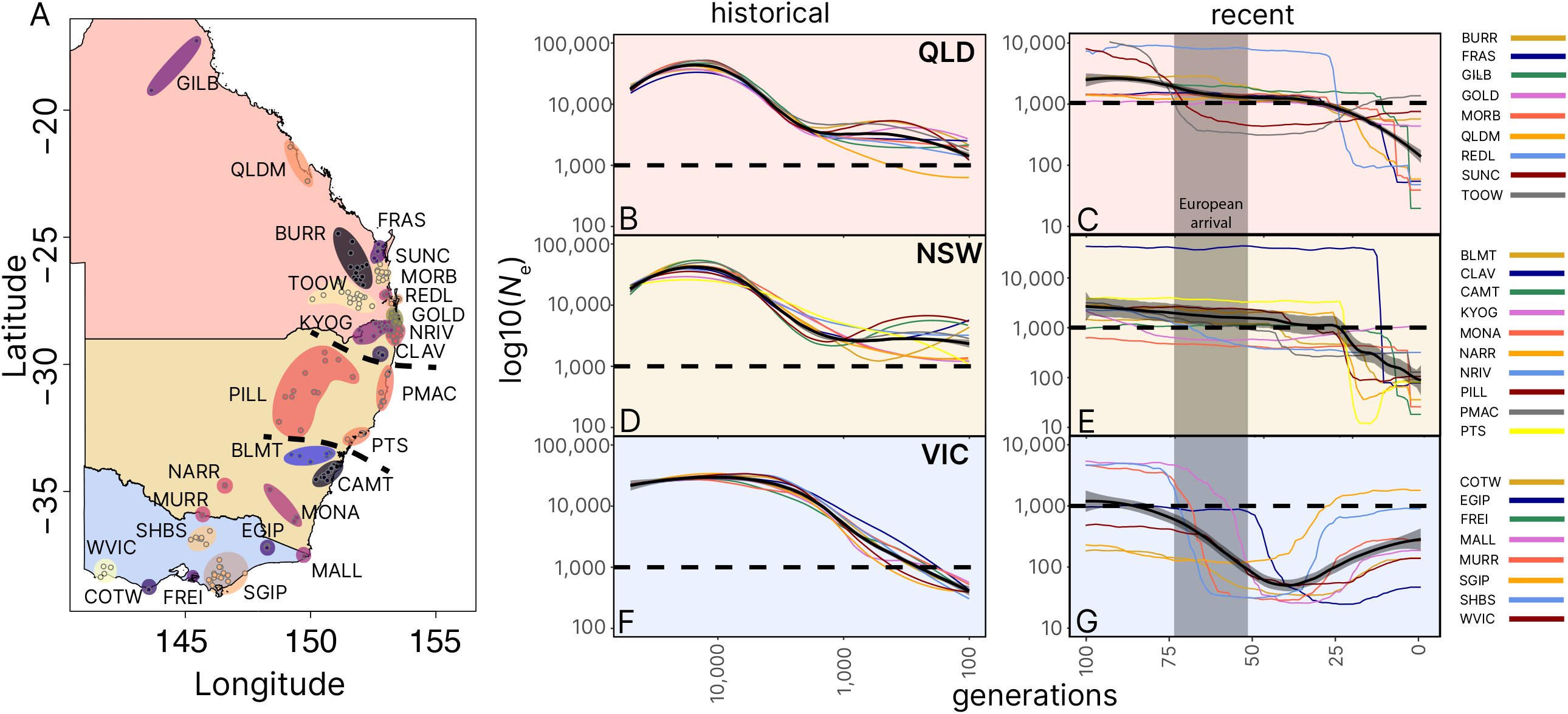
Distribution and demographic histories of koala populations used in the WGS analyses. (A) Spatial map of the 27 populations used in the study. (B, D, F) Reconstruction of historical demographic history from SMC++ for each koala population segregated into different geopolitical states (VIC = Victoria, NSW = New South Wales, QLD = Queensland). (C, E, G) Reconstruction of recent *N*_e_ estimates from GONE for each population. The black line in each figure represents the mean over populations and the shaded areas are 99% confidence intervals.

Recent changes in *N*_e_ (*21*) revealed stark contrasts among state meta-populations. Our analyses indicate the recovery of the bottlenecked VIC meta-population started approximately 40 generations ago (Fig. 1J), whereas populations in QLD and NSW show sharp declines around 10-30 generations ago (Fig. 1DG). Within VIC, the SGIP population displayed the highest final *N*_e_ among all studied populations (Fig. 1; all population codes are spatially defined in Fig. 1A and specified in Table S1), corroborated by *N*_e_ inferred from runs of homozygosity (ROH) (Fig. S1). Some QLD and NSW populations appear to have plateaued or show slight *N*_e_ increases, while most others, such as CLAV and CAMT in NSW, and MORB and FRAS in QLD, exhibit recent declines in *N*_e_.

Critically, these demographic reconstructions align with historical management records. The catastrophic decline experienced in VIC in the late 1800s is clear from our historical reconstruction of *N*_e_, as is a rapid increase in *N*_e_ for most VIC populations, which corresponds with their reintroduction histories. In particular, the COTW population, founded in 1981, grew to a population size of >10,000 by 2013 and was the target of intensive management (mass culling, translocations and fertility control) during 2013-18 to reduce welfare issues from over-browsing (*15*). Current *N*_e_ estimates from LD suggest that the COTW population has the highest *N*_e_, more than twice that of any other population (Table S1). This likely reflects these recent population dynamics, where the population explosion has led to more recombination, rather than an increase in rare alleles (see below).

### Genetic diversity and inbreeding

Our results indicate that patterns of genetic diversity and inbreeding within koala populations may be more complex than currently assumed. Previous studies have reported that VIC populations exhibit significantly lower genetic diversity compared to those from NSW and QLD (*22,23*). While state-level patterns remain broadly consistent (p < 0.001; Fig. 2A), the degree of differentiation is less pronounced when compared to prior studies that filter datasets by minor allele frequency (MAF) and LD. Previous findings reported NSW and QLD populations having 1.76 times the heterozygosity of VIC, whereas our analysis, which includes rare alleles, indicates a ratio of only 1.30.

**Figure 2.**
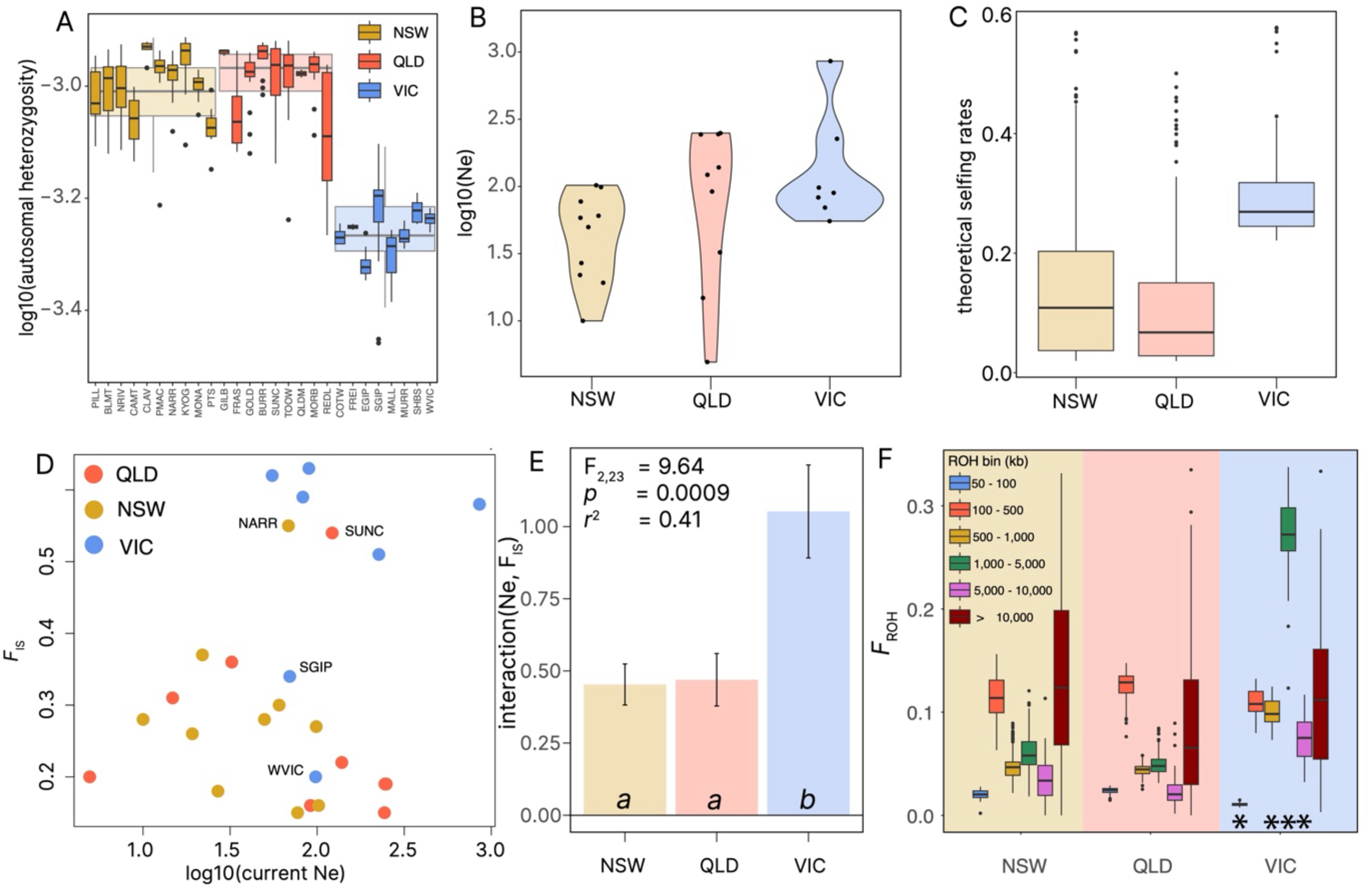
Inbreeding and diversity summary statistics across all 27 populations. (A) Population boxplots for autosomal heterozygosity and their state-of-origin (color) with significance between states identified at an α level of 0.05. (B) Current effective population size (*N*_e_) estimated using currentNe for each population. (C) Theoretical selfing rates for each population determined by runs of homozygosity (ROH), Tajima’s D, and F statistics using roh-selfing. (D) Scatter plot of the relationship between population level inbreeding coefficients (*F*) and current measures of *N*_e_. (E) Differences between the interaction of *F* and *N*_e_ between states with significance at the α level of 0.05. (F) Inbreeding estimated by runs of homozygosity (*F*_ROH_) split into six ROH length bins. Significant differences between Victoria and other states denoted by * at an α = 0.05.

Estimates of contemporary *N*_e_ challenge the notion of VIC populations being uniformly devoid of genetic diversity (Fig. 2B; *p* = 0.16) and contrast with theoretical selfing-rates (Fig. 2C; *F*_2,294_ = 46.39; p < 0.001). Additionally, individual inbreeding (*F*) does not appear to correlate directly with contemporary *N*_e_ (Fig. 2D; *F*_1,24_ = 0.76; p = 0.39). Instead, the interaction between *F* and contemporary *N*_e_ across states reveals significant differences where populations with the highest *F* also have the greatest *N*_e_ (Fig. 2E; *F*_2,23_ = 9.64; p < 0.001; r^2^ = 0.41).

Further insights emerge from analyses of ROH, which show that VIC populations have fewer small ROHs (reflective of past bottlenecks) but a greater prevalence of intermediate-length ROHs (reflective of recent bottlenecks) compared to QLD and NSW (Fig. 2F). Cumulative inbreeding measures based on ROHs (*F*_ROH_) are more accurate than other measures (*24*) and highlight state-level differences, even though our data shows that the individual measures of *F*_ROH_ and *F* are highly correlated (Fig. S2). Specifically, koalas from VIC have on average 68.1% of their genome covered by ROHs >50 kb, compared to 41.2% in NSW and 36.0% in QLD. Although long ROHs (>10,000 kb) do not differ significantly between states (p = 0.42 and p = 0.06, respectively), intermediate ROHs (>500 and <10,000 kb) are significantly greater in VIC (all t-values > 7; all *p* < 0.001; Fig. 2F and Fig. S3). VIC has a significantly smaller *F*_ROH_ in the shortest ROH bin (>50 and <100 kb; t-value = -8.34; p < 0.001). Significant differences in the shortest and intermediate ROH lengths suggest that VIC populations experienced significant bottlenecks or founder events at several time points (*25*), consistent with the trends reflected in our demographic reconstructions and known demographic characteristics.

### Distribution of alleles

While patterns of genetic diversity provide valuable insights into past demographic histories, they do not fully explain these recent trends in *N*_e_. Demographic expansion is expected to result in more recombination events within populations and increased mutation leading to the accumulation of rare alleles. Rare alleles have been shown to play crucial roles in human biology, having a significant influence on susceptibility to disease (*26*) and adaptation to novel environments (*27*). We therefore hypothesized that there may be an increase in rare alleles in populations that have undergone demographic expansion (increased *N*_e_), while acknowledging the recency of expansion events and overall population sizes as key influential factors. To test this hypothesis, we partitioned autosomal heterozygosity into three discrete minor allele frequency (MAF) bins: 1) common alleles (MAF > 0.05), 2) low frequency alleles (MAF 0.05–0.01), and 3) rare alleles (MAF < 0.01).

The common allele group exhibited significant state-level differences that were consistent with previous analyses (*23*), with VIC populations displaying the lowest diversity and QLD populations the highest (Fig. 3A; *F*_3,426_ = 53.17; p < 0.001), while intra-state variation was relatively consistent (Fig. 3A). For the low frequency allele group, a similar overall pattern was observed (*F*_3,426_ = 329.8; p < 0.001), although VIC populations displayed greater variability (Fig. 3B). In the rare allele group, VIC showed statistically lower heterozygosity than NSW, though considerable interstate population overlap was evident (Fig. 3C; *F*_3,426_ = 228.6; p < 0.001). We then explored the ratio between MAF bins to determine if rare alleles are being recovered in VIC populations based on the theory that an allele frequency is correlated with its probability of being lost during a bottleneck (*28, 29*). Critically, we found major differences between VIC and other states in the low/common ratio (Fig. 3D) as expected, but fewer differences between states in the rare/low ratio, especially for the SGIP, SHBS, and WVIC populations (Fig. 3E). This dynamic suggests that genetic recovery has begun through novel mutations in the rare MAF bin.

**Figure 3.**
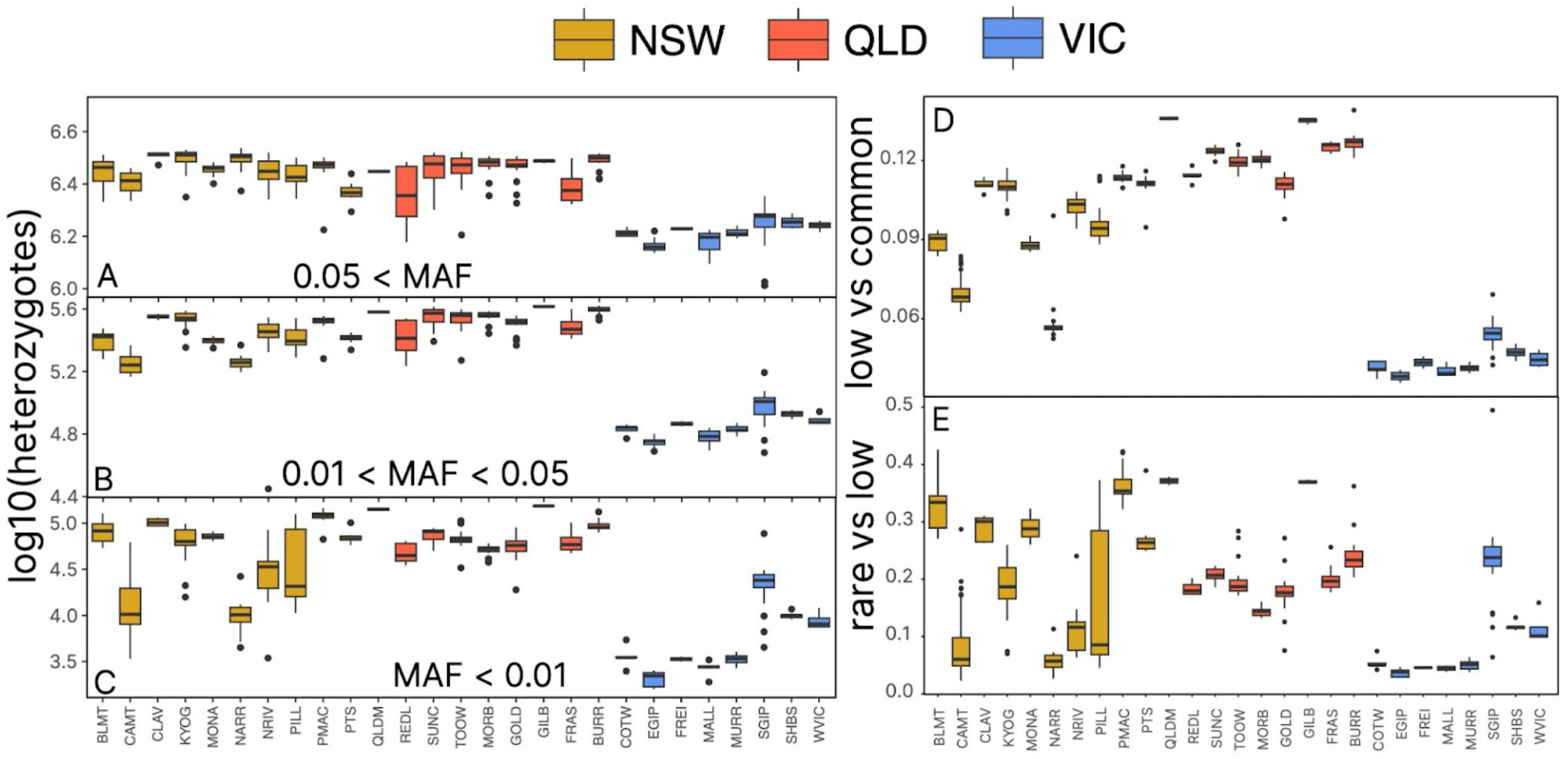
Distribution of heterozygotes across populations in three different allele frequency bins. (A,B,C) Boxplots of heterozygosity for the three minor allele frequency (MAF) bins for each population, where (A) is MAF >0.05, (B) is MAF 0.01 - 0.05, (C) is MAF <0.01. (D,E) Boxplots describing the ratio of autosomal heterozygotes between MAF bin pairs, with (D) being MAF_LOW_ /MAF_COM_ and (E) is the MAF_RARE_ /MAF_LOW_.

Principle component analysis also illustrates how population structure varies with allele frequency bins, transitioning from regional differentiation at common allele frequencies (state-based; Fig. S4A,B), to local differentiation at low allele frequencies (population-based; Fig. S4C,D), and finally to individual-level variation at rare allele frequencies (Fig. S4E,F). Variability between populations increased as the MAF threshold decreased, with two NSW populations (CAMT and NARR) showing pronounced declines in heterozygosity across allele frequency bins (Fig. 3C). For NARR, this decline likely reflects genetic drift in an isolated population that is known to be derived from two introduced divergent parental populations (*30*). Previous work suggests that this population has comparable genetic diversity to other southern NSW populations (*23*); however, it is diverse for common alleles only. In contrast, it has the lowest number of heterozygotes in both low and rare frequency allele bins among all NSW and QLD populations, suggesting rare alleles were lost due to low population size and associated drift processes.

Within VIC, populations such as SGIP, SHBS, and WVIC exhibited higher heterozygosity than other VIC populations, particularly within the low and rare allele bins (Fig. 3D, E). The SGIP and SHBS populations are thought to contain remnant genetic variants (*31*), despite the introduction of French Island individuals into these regions (*15*). In contrast, the WVIC population was reintroduced in the 1970s (*15*), experienced rapid demographic growth within a few decades, and subsequently required management intervention (culling, relocation, sterilisation). The elevated heterozygosity observed in WVIC is therefore likely attributable to mutation during this period of sustained population expansion, particularly given its earlier recovery relative to populations such as COTW. These findings highlight previously unrecognised heterogeneity within the VIC metapopulation that may be important for the species’ long-term evolutionary potential. Notably, this contrasts with earlier assessments that broadly characterized Victorian koalas as genetically depauperate (*22,23*).

### Functional genetic diversity

Genetic load is complex, often environment-dependent, generally interpretable from genomic data only when accompanying ecological and phenotypic information, and requires experimental validation (*32*). Although genetic load can negatively impact fitness through the accumulation of deleterious alleles, purging of deleterious alleles can occur in severely bottlenecked populations and reduce the fitness costs of inbreeding (*33*). Because it is impossible to know if loss-of-function (LOF) variants are deleterious (*34–38*), we consider consequential variation in several functional groups (LOF, regulatory, and missense mutations) across populations.

We used the *R*_X/Y_ metric (*39*), a pairwise statistic used to compare the frequency of mutations between two populations and to estimate the effects of purifying selection, to assess whether demographic expansion has influenced the accumulation of functional variants. As predicted by theory, which suggests that population growth can lead to an accumulation of functional variants (*40*), VIC populations were found to carry a higher proportion of LOF, regulatory and missense variants across all MAF bins (Fig. 4A–C). This pattern is consistent with a recent expansion from a small founder population, in which relaxed selection and increased mutational input have contributed to the accumulation of functional variants.

**Figure 4.**
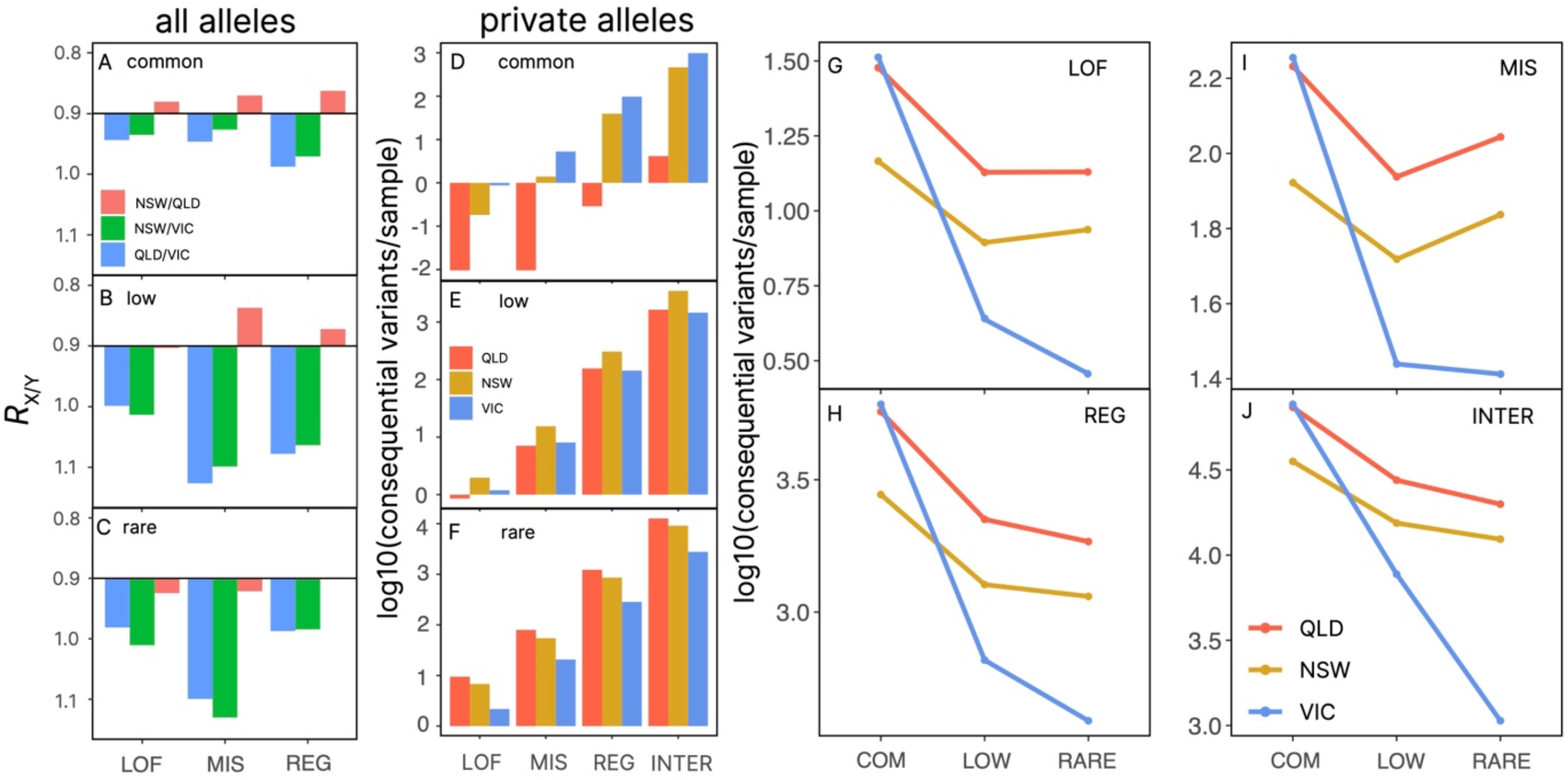
Differences between functional variation within and between populations. (A,B,C) Pairwise *R*_X/Y_ estimates between Queensland (QLD), New South Wales (NSW) and Victoria (VIC). Interpretation is that an *R*_X/Y_ > 1 indicates that population X (numerator) has a higher rate of functional alleles than Y, and *R*_X/Y_ < 1 indicates that population Y (denominator) has a higher rate of functional alleles. (D,E,F) State changes of private functional alleles across MAF bins for each states. (G,H,I,J) Changes in the number of variants per sample for each variant type across MAF bins for each state. Without a bottleneck, our expectations are met with QLD and NSW meta-populations, similar findings for all bins and variant types. On the other hand, the VIC meta-populations also meet our expectations with the number of low-frequency functional variants being much lower than the common bin (suggestive of a bottleneck), while the number of variants in the rare bin remains similar to the low frequency bin (suggestive of an expansion).

To disentangle the effects state specific functional variation, we further partitioned variants by state and MAF bin assignment. When examining private alleles per sample, VIC was enriched for functional variants in the common frequency bin (Fig. 4D), NSW in the low-frequency bin (Fig. 4E), and QLD in the rare-frequency bin (Fig. 4F). LOF variants were most frequent in low-frequency bins, consistent with theoretical expectations for purifying selection acting on deleterious alleles. Each state possesses many private variants, suggesting that the less diverse VIC meta-population still retains substantial variation not found elsewhere and may be important for future adaptation.

The influence of demographic history was further evident in total variant counts per individual (Fig. 4G-J). NSW had fewer common functional and intergenic variants than VIC and QLD, while VIC had the fewest rare and low-frequency variants—consistent with a severely bottlenecked population. Interestingly, VIC lost proportionally more intergenic than functional variants, suggesting that different evolutionary forces may be acting on neutral versus functional genomic regions within these populations and explains the intrastate patterns from the *R*_X/Y_ analysis. Additionally, VIC populations have lost more low and rare frequency functional variants than expected when compared to patterns from other states, suggestive of purging in the VIC populations.

Together, these findings indicate that each koala population maintains a distinct spectrum of low- and rare-frequency functional alleles, reflecting their unique demographic histories and evolutionary trajectories. The divergence among states underscores the importance of considering both shared and private variation when evaluating adaptive potential in wildlife genomics and conservation planning.

## Discussion

Our findings challenge the prevailing assumption that genetic bottlenecks in threatened species constrain long-term population viability through irreversible loss of genetic variation and accumulation of inbreeding load (*41*). Despite a well-documented bottleneck and strikingly low genetic diversity (*16,23,42*), koalas in Victoria exhibit signatures of demographic recovery evidenced by an increasing *N*_e_ and functional variation that is comparable, and in some cases greater, relative to other states. These results parallel the *genetic paradox of invasive species* (*43*), where natural populations can ‘escape’ inbreeding following extreme bottlenecks. Our results demonstrate that demographic expansion can play a critical role in reshaping the evolutionary trajectory of bottlenecked threatened species.

We argue that recombination plays a central role in the early stages of recovery following a population bottleneck. In expanding koala populations, increased recombination appears to have accelerated linkage disequilibrium (LD) decay (Fig. S5) and facilitated the reassortment of genetic variation, consistent with observed increases in *N*_e_. Despite its importance, the role of recombination in enhancing population viability is often underappreciated in conservation biology. Recombination can disrupt deleterious allele combinations (*44*), enhance adaptive potential (*12,45*), and accelerate evolutionary responses to selection (*12,46*). These benefits arise from the reassortment of genetic variation among individuals, which increases the range of trait combinations available to selection (*12*). As a result, traditional metrics of genetic diversity may not fully capture a population’s evolutionary capacity; what matters is not only the number of alleles present, but how they are recombined. This helps explain why we observe cases of low genetic diversity co-occurring with high *N*_e_, and vice versa.

To our knowledge, this is the first time that both patterns have been identified in different populations of the same species. However, low diversity with constant *N*_e_ was found in New Zealand snapper (*Chrysophrys auratus*) (*47*), and the opposite was identified with temporally collected kit fox samples (*48*).

While overall genetic diversity remains low in southern koala populations, we observed a surprising abundance of low-frequency, private and functional variants—especially in populations with longer histories of demographic expansion. This suggests that *de novo* mutations, recombination, and relaxed selection contribute to the accumulation of novel variation across the genome and its differential assortment, even in the wake of extreme bottlenecks. It also suggests that the processes of purging and accumulation of rare variation are ephemeral and contingent on demographic contraction and expansion, respectively. These dynamics offer a mechanism by which small, threatened populations “escape” inbreeding, not through restoring ancestral diversity, but by generating new alleles and purging mildly deleterious alleles while increasing the efficiency of selection via recombination. Theoretically, these evolutionary processes could create healthier populations with fewer deleterious alleles and greater combinations of novel variation (*10,49*). Currently, Victorian koala populations are on this trajectory, which contrasts with their northern counterparts in New South Wales and Queensland that have declining *N*_e_, raising concerns about ongoing genetic erosion, particularly as habitat fragmentation, disease, and climatic stressors continue to aXect populations in these regions.

While Victorian populations are amidst a remarkable demographic recovery and now carry increasing numbers of novel, reassorted and potentially adaptive variants, their genetic architecture remains vulnerable. A future bottleneck—caused by disease, habitat loss, human interventions, or extreme climate events—could easily erase this newly accumulated diversity. Management strategies must therefore be forward-looking, recognizing the importance of population expansion and preserving rare alleles across populations. Consideration should also be given to further increasing the resilience of the species through genetic mixing between Victorian and New South Wales or Queensland populations, which may also help to safeguard the genetic variation found in northern declining populations (*50,51*). We advise that a national strategy should prioritize the preservation and connectivity of koala populations to enhance their evolutionary potential and mitigate ongoing threats.

Taken together, our results demonstrate that demographic history interacts with recombination, mutation and human intervention in complex ways to shape contemporary patterns of genetic variation. Far from being uniformly depleted, bottlenecked populations may retain or even regenerate functional variation, particularly during demographic recovery. This has profound implications for conservation genomics: (i) low genetic diversity does not necessarily imply limited evolutionary potential, (ii) inbreeding and diversity metrics are context-dependant and shaped by demographic history, and (iii) management strategies must consider both recent population trajectories and the distribution of rare and functional alleles. The koala provides a compelling example of how, under the right demographic conditions, a severely bottlenecked species can escape the genetic consequences of inbreeding and recover through natural evolutionary processes.

## Acknowledgements

We acknowledge the Koala Genome Survey, NSW Government, and the Australian Government’s Bushfire Recovery for Wildlife for their roles in generating the publicly available data.

## References

1. Y. Willi, T. N. Kristensen, C.M. Sgrò, A. R. Weeks, M. Ørsted, A. A. Hoffmann, Conservation genetics as a management tool: The five best-supported paradigms to assist the management of threatened species. Proc National Acad Sci 119, e2105076119 (2022).

2. T. Madsen, B. Stille, R. Shine, Inbreeding depression in an isolated population of adders Vipera berus. Biol. Conserv. 75, 113–118 (1996).

3. D. Charlesworth, J. H. Willis, The genetics of inbreeding depression. Nat Rev Genet 10, 783–796 (2009).

4. D. A. Roff, Evolutionary Quantitative Genetics (Chapman & Hall, 1997)

5. L. F. Keller, D. M. Waller, Inbreeding effects in wild populations. Trends Ecol Evol 17, 230–241 (2002).

6. S. J. O’Brien, M. E. Roelke, L. Marker, A. Newman, C. A. Winkler, D. Meltzer, L. Colly, J. F. Evermann, M. Bush, D. E. Wildt, Genetic Basis for Species Vulnerability in the Cheetah. Science 227, 1428–1434 (1985).

7. Y. Willi, J. V. Buskirk, A. A. Hoffmann, Limits to the Adaptive Potential of Small Populations. Annu Rev Ecol Evol Syst 37, 433–458 (2006).

8. A. A. Hoffmann, C.M. Sgrò, T. N. Kristensen, Revisiting Adaptive Potential, Population Size, and Conservation. Trends Ecol Evol 32, 506–517 (2017).

9. D. Blomqvist, A. Pauliny, M. Larsson, L.-Å. Flodin, Trapped in the extinction vortex? Strong genetic effects in a declining vertebrate population. BMC Evol. Biol. 10, 33 (2010).

10. A. Khan, K. Patel, H. Shukla, A. Viswanathan, T. van der Valk, U. Borthakur, P. Nigam, A. Zachariah, Y. V. Jhala, M. Kardos, U. Ramakrishnan, Genomic evidence for inbreeding depression and purging of deleterious genetic variation in Indian tigers. Proc. Natl. Acad. Sci. 118, e2023018118 (2021).

11. S. Wright, Evolution in mendelian populations. Genetics 16, 97–159 (1931).

12. J. V. Peñalba, J. B. W. Wolf, From molecules to populations: appreciating and estimating recombination rate variation. Nat. Rev. Genet. 21, 476–492 (2020).

13. R. Lande, Genetics and Demography in Biological Conservation. Science 241, 1455– 1460 (1988).

14. A. Estoup, V. Ravigné, R. Hufbauer, R. Vitalis, M. Gautier, B. Facon, Is There a Genetic Paradox of Biological Invasion? Annu. Rev. Ecol., Evol., Syst. 47, 1–22 (2016).

15. P. Menkhorst, Hunted, marooned, re-introduced, contracepted: a history of koala management in Victoria. in Too close for comfort: contentious issues in human-wildlife encounters. Royal Zoological Society of New South Wales, Mosman, NSW, Australia. 73–92 (2008).

16. B. A. Houlden, P. R. England, A. C. Taylor, W. D. Greville, W. B. Sherwin, Low genetic variability of the koala Phascolarctos cinereus in south-eastern Australia following a severe population bottleneck. Mol. Ecol. 5, 269–281 (1996).

17. G. Heard, D. Ramsey, Modelling koala abundance across Victoria. unpublished client report (2020).

18. C. J. Hogg, L. Silver, E. A. McLennan, K. Belov, Koala Genome Survey: An Open Data Resource to Improve Conservation Planning. Genes-basel 14, 546 (2023).

19. J. Terhorst, J. A. Kamm, Y. S. Song, Robust and scalable inference of population history from hundreds of unphased whole genomes. Nat. Genet. 49, 303–309 (2017).

20. R. N. Johnson, D. O’Meally, Z. Chen, G. J. Etherington, S. Y. W. Ho, W. J. Nash, C. E. Grueber, Y. Cheng, C. M. Whittington, S. Dennison, E. Peel, W. Haerty, R. J. O’Neill, D. Colgan, T. L. Russell, D. E. Alquezar-Planas, V. Attenbrow, J. G. Bragg, P. A. Brandies, A. Y.-Y. Chong, J. E. Deakin, F. D. Palma, Z. Duda, M. D. B. Eldridge, K. M. Ewart, C. J. Hogg, G. J. Frankham, A. Georges, A. K. Gillett, M. Govendir, A. D. Greenwood, T. Hayakawa, K. M. Helgen, M. Hobbs, C. E. Holleley, T. N. Heider, E. A. Jones, A. King, D. Madden, J. A. M. Graves, K. M. Morris, L. E. Neaves, H. R. Patel, A. Polkinghorne, M. B. Renfree, C. Robin, R. Salinas, K. Tsangaras, P. D. Waters, S. A. Waters, B. Wright, M. R. Wilkins, P. Timms, K. Belov, Adaptation and conservation insights from the koala genome. Nat. Genet. 50, 1102–1111 (2018).

21. E. Santiago, I. Novo, A. F. Pardiñas, M. Saura, J. Wang, A. Caballero, Recent Demographic History Inferred by High-Resolution Analysis of Linkage Disequilibrium. Mol. Biol. Evol. 37, 3642–3653 (2020).

22. M. J. Lott, B. R. Wright, L. E. Neaves, G. J. Frankham, S. Dennison, M. D. B. Eldridge, S. Potter, D. E. Alquezar-Planas, C. J. Hogg, K. Belov, R. N. Johnson, Future-proofing the koala: Synergising genomic and environmental data for effective species management. Mol. Ecol. 31, 3035–3055 (2022).

23. E. A. McLennan, T. G. L. Kovacs, L. W. Silver, Z. Chen, F. R. Jaya, S. Y. W. Ho, K. Belov, C. J. Hogg, Genomics identifies koala populations at risk across eastern Australia. Ecol. Appl., e3062 (2024).

24. A. Caballero, B. Villanueva, T. Druet, On the estimation of inbreeding depression using different measures of inbreeding from molecular markers. Evol. Appl. 14, 416– 428 (2021).

25. A. D. Foote, R. Hooper, A. Alexander, R. W. Baird, C. S. Baker, L. Ballance, J. Barlow, A. Brownlow, T. Collins, R. Constantine, L. D. Rosa, N. J. Davison, J. W. Durban, R. Esteban, L. Excoffier, S. L. F. Martin, K. A. Forney, T. Gerrodette, M. T. P. Gilbert, C. Guinet, M. B. Hanson, S. Li, M. D. Martin, K. M. Robertson, F. I. P. Samarra, R. de Stephanis, S. B. Tavares, P. Tixier, J. A. Totterdell, P. Wade, J. B. W. Wolf, G. Fan, Y. Zhang, P. A. Morin, Runs of homozygosity in killer whale genomes provide a global record of demographic histories. Mol. Ecol. 30, 6162–6177 (2021).

26. R. D. Hernandez, L. H. Uricchio, K. Hartman, C. Ye, A. Dahl, N. Zaitlen, Ultrarare variants drive substantial cis heritability of human gene expression. Nat Genet 51, 1349–1355 (2019).

27. D. Agashe, M. Sane, S. Singhal, Revisiting the Role of Genetic Variation in Adaptation. Am. Nat. 202, 486–502 (2023).

28. W. Amos, J. I. Hoffman, Evidence that two main bottleneck events shaped modern human genetic diversity. Proc. R. Soc. B: Biol. Sci. 277, 131–137 (2009).

29. F. W. Allendorf, O. Hössjer, N. Ryman, What does effective population size tell us about loss of allelic variation? Evol. Appl. 17, e13733 (2024).

30. J. A. Sullivan, “Brief History of Koala Regeneration Centre.” (1990); Retrieved from Narranderal Koala Regeneration Centre Super-visory Committee.

31. F. Wedrowicz, J. Mosse, W. Wright, F. E. Hogan, Genetic structure and diversity of the koala population in South Gippsland, Victoria: a remnant population of high conservation significance. Conserv. Genet. 19, 713–728 (2018).

32. G. Bertorelle, F. Raffini, M. Bosse, C. Bortoluzzi, A. Iannucci, E. Trucchi, H. E. Morales, C. van Oosterhout, Genetic load: genomic estimates and applications in non-model animals. Nat. Rev. Genet. 23, 492–503 (2022).

33. P. W. Hedrick, A. Garcia-Dorado, Understanding Inbreeding Depression, Purging, and Genetic Rescue. Trends Ecol. Evol. 31, 940–952 (2016).

34. D. G. MacArthur, S. Balasubramanian, A. Frankish, N. Huang, J. Morris, K. Walter, L. Jostins, L. Habegger, J. K. Pickrell, S. B. Montgomery, C. A. Albers, Z. D. Zhang, D. F. Conrad, G. Lunter, H. Zheng, Q. Ayub, M. A. DePristo, E. Banks, M. Hu, R. E. Handsaker, J. A. Rosenfeld, M. Fromer, M. Jin, X. J. Mu, E. Khurana, K. Ye, M. Kay, G. I. Saunders, M.-M. Suner, T. Hunt, I. H. A. Barnes, C. Amid, D. R. Carvalho-Silva, A. H. Bignell, C. Snow, B. Yngvadottir, S. Bumpstead, D. N. Cooper, Y. Xue, I. G. Romero, 1000 Genomes Project Consortium, J. Wang, Y. Li, R. A. Gibbs, S. A. McCarroll, E. T. Dermitzakis, J. K. Pritchard, J. C. Barrett, J. Harrow, M. E. Hurles, M. B. Gerstein, C. Tyler-Smith, A Systematic Survey of Loss-of-Function Variants in Human Protein-Coding Genes. Science 335, 823–828 (2012).

35. C. L. Shovlin, M. A. Aldred, When “loss-of-function” means proteostasis burden: Thinking again about coding DNA variants. Am. J. Hum. Genet. 112, 3–10 (2025).

36. J. Serrano, S. Kondo, G. M. Link, I. S. Brown, R. E. Pratley, K. K. Baskin, B. H. Goodpaster, P. M. Coen, G. A. Kyriazis, A partial loss-of-function variant (Ile191Val) of the TAS1R2 glucose receptor is associated with enhanced responses to exercise training in older adults with obesity: A translational study. Metabolism 162, 156045 (2025).

37. N. Whiffin, K. J. Karczewski, X. Zhang, S. Chothani, M. J. Smith, D. G. Evans, A. M. Roberts, N. M. Quaife, S. Schafer, O. Rackham, J. Alföldi, A.H. O’Donnell-Luria, L. C. Francioli, I. M. Armean, E. Banks, L. Bergelson, K. Cibulskis, R. L. Collins, K. M. Connolly, M. Covarrubias, B. Cummings, M. J. Daly, S. Donnelly, Y. Farjoun, S. Ferriera, S. Gabriel, L. D. Gauthier, J. Gentry, N. Gupta, T. Jeandet, D. Kaplan, K. M. Laricchia, C. Llanwarne, E. V. Minikel, R. Munshi, B. M. Neale, S. Novod, N. Petrillo, T. Poterba, D. Roazen, V. Ruano-Rubio, A. Saltzman, K. E. Samocha, M. Schleicher, C. Seed, M. Solomonson, J. Soto, G. Tiao, K. Tibbetts, C. Tolonen, C. Vittal, G. Wade, A. Wang, Q. Wang, N. A. Watts, B. Weisburd, C. A. A. Salinas, T. Ahmad, C. M. Albert, D. Ardissino, G. Atzmon, J. Barnard, L. Beaugerie, E. J. Benjamin, M. Boehnke, L. L. Bonnycastle, E. P. Bottinger, D. W. Bowden, M. J. Bown, J. C. Chambers, J. C. Chan, D. Chasman, J. Cho, M. K. Chung, B. Cohen, A. Correa, D. Dabelea, M. J. Daly, D. Darbar, R. Duggirala, J. Dupuis, P. T. Ellinor, R. Elosua, J. Erdmann, T. Esko, M. Färkkilä, J. Florez, A. Franke, G. Getz, B. Glaser, S. J. Glatt, D. Goldstein, C. Gonzalez, L. Groop, C. Haiman, C. Hanis, M. Harms, M. Hiltunen, M. M. Holi, C. M. Hultman, M. Kallela, J. Kaprio, S. Kathiresan, B.-J. Kim, Y. J. Kim, G. Kirov, J. Kooner, S. Koskinen, H. M. Krumholz, S. Kugathasan, S. H. Kwak, M. Laakso, T. Lehtimäki, R. J. F. Loos, S. A. Lubitz, R. C. W. Ma, J. Marrugat, K. M. Mattila, S. McCarroll, M. I. McCarthy, D. McGovern, R. McPherson, J. B. Meigs, O. Melander, A. Metspalu, B. M. Neale, P. M. Nilsson, M. C. O’Donovan, D. Ongur, L. Orozco, M. J. Owen, C. N. A. Palmer, A. Palotie, K. S. Park, C. Pato, A. E. Pulver, N. Rahman, A. M. Remes, J. D. Rioux, S. Ripatti, D. M. Roden, D. Saleheen, V. Salomaa, N. J. Samani, J. Scharf, H. Schunkert, M. B. Shoemaker, P. Sklar, H. Soininen, H. Sokol, T. Spector, P. F. Sullivan, J. Suvisaari, E. S. Tai, Y. Y. Teo, T. Tiinamaija, M. Tsuang, D. Turner, T. Tusie-Luna, E. Vartiainen, H. Watkins, R. K. Weersma, M. Wessman, J. G. Wilson, R. J. Xavier, M. P. Vawter, S. A. Cook, P. J. R. Barton, D. G. MacArthur, J. S. Ware, Characterising the loss-of-function impact of 5’ untranslated region variants in 15,708 individuals. Nat. Commun. 11, 2523 (2020).

38. Y.-C. Xu, Y.-L. Guo, Less Is More, Natural Loss-of-Function Mutation Is a Strategy for Adaptation. Plant Commun. 1, 100103 (2020).

39. R. Do, D. Balick, H. Li, I. Adzhubei, S. Sunyaev, D. Reich, No evidence that selection has been less effective at removing deleterious mutations in Europeans than in Africans. Nat. Genet. 47, 126–131 (2015).

40. J. A. Tennessen, A. W. Bigham, T. D. O’Connor, W. Fu, E. E. Kenny, S. Gravel, S. McGee, R. Do, X. Liu, G. Jun, H. M. Kang, D. Jordan, S. M. Leal, S. Gabriel, M. J. Rieder, G. Abecasis, D. Altshuler, D. A. Nickerson, E. Boerwinkle, S. Sunyaev, C. D. Bustamante, M. J. Bamshad, J. M. Akey, B. GO, S. GO, on behalf of the N. E. S. Project, Evolution and Functional Impact of Rare Coding Variation from Deep Sequencing of Human Exomes. Science 337, 64–69 (2012).

41. P. Nietlisbach, S. Muff, J. M. Reid, M. C. Whitlock, L. F. Keller, Nonequivalent lethal equivalents: Models and inbreeding metrics for unbiased estimation of inbreeding load. Evol. Appl. 12, 266–279 (2019).

42. A. C. Taylor, J. M. Graves, N. D. Murray, S. J. O’brien, N. Yuhki, B. Sherwin, Conservation genetics of the koala (Phascolarctos cinereus): low mitochondrial DNA variation amongst southern Australian populations. Genet. Res. 69, 25–33 (1997).

43. R. I. Colautti, J. M. Alexander, K. M. Dlugosch, S. R. Keller, S. E. Sultan, Invasions and extinctions through the looking glass of evolutionary ecology. Philos. Trans. R. Soc. B: Biol. Sci. 372, 20160031 (2017).

44. N. H. Barton, Mutation and the evolution of recombination. Philos. Trans. R. Soc. B: Biol. Sci. 365, 1281–1294 (2010).

45. S. P. Otto, N. H. Barton, Selection for recombination in small populations. Evolution 55, 1921–1931 (2001).

46. T. F. Cooper, Recombination Speeds Adaptation by Reducing Competition between Beneficial Mutations in Populations of Escherichia coli. PLoS Biol. 5, e225 (2007).

47. L. Hauser, G. J. Adcock, P. J. Smith, J. H. B. Ramírez, G. R. Carvalho, Loss of microsatellite diversity and low effective population size in an overexploited population of New Zealand snapper (Pagrus auratus). Proc. Natl. Acad. Sci. 99, 11742–11747 (2002).

48. R. C. Lonsinger, J. R. Adams, L. P. Waits, Evaluating effective population size and genetic diversity of a declining kit fox population using contemporary and historical specimens. Ecol. Evol. 8, 12011–12021 (2018).

49. A. Caballero, I. Bravo, J. Wang, Inbreeding load and purging: implications for the short-term survival and the conservation management of small populations. Heredity 118, 177–185 (2017).

50. A. R. Weeks, C. M. Sgro, A. G. Young, R. Frankham, N. J. Mitchell, K. A. Miller, M. Byrne, D. J. Coates, M. D. B. Eldridge, P. Sunnucks, M. F. Breed, E. A. James, A. A. Hoffmann, Assessing the benefits and risks of translocations in changing environments: a genetic perspective. Evol Appl 4, 709–725 (2011).

51. A. A. Hoffmann, A. D. Miller, A. R. Weeks, Genetic mixing for population management: From genetic rescue to provenancing. Evol Appl 14, 634–652 (2021).

