## Supplementary Information for "Escaping inbreeding: the demographic path to genetic recovery"

### Methods

**Samples and bioinformatics**

This study made use of publicly available whole genome resequencing data for 418 koala samples from 27 spatially defined populations spanning the species' entire distribution on mainland Australia (Table S1; *1*) . Sequence data from each sample was aligned to a publicly available koala genome (NCBI ID: GCA_003287225.2) using the Illumina DRAGEN pipeline, an efficient and accurate variant caller (*2*), which generated a gvcf file for each sample. Variant calling was performed using the Joint Genotyper pipeline implemented in DRAGEN, which uses a GATK algorithm to improve accuracy and processing times (*3*), and creates VCF files for downstream analyses. Joint Genotyper only accepts 99 files at a time, therefore we used the pipeline to process the data in five separate batches and subsequently merged all vcf files using bcftools. Variants that were missing from a batch were considered homozygous for the reference allele. Because our research questions revolved around assessments of runs of homozygosity (ROH), it was imperative that we focused on long regions, and therefore dataset was reduced to the 11 longest scaffolds (accounting for 98.9% of the assembled genome), henceforth called pseudo chromosomes. This initial full vcf file had ~46.8 million single nucleotide variants (SNVs). The variant file was then filtered by read depth (>10 and <100 read depth; to avoids issues of associated with paralogy), quality (>30 phred), and limited to biallelic SNVs, using bcftools. These thresholds were used for all datasets, but several different additional filtering methods were implemented (e.g., missing data, minor allele frequency and minor allele count) to optimise the dataset for specific analyses (Table S2 shows the filtering of each dataset). After filtering, the base dataset consisted of 26,307,311 single nucleotide polymorphisms (SNPs).

#### Analyses

Demographic history reconstruction: We use three methods to reconstruct demographic history, including reconstructions of both historical to recent effective population sizes ($N_e$). Each analysis was performed for each of the 27 defined populations, but also involved the analysis of populations by state for visualisation purposes, due to (1) each state having different histories of hunting/culling, (2) each state having different management histories and strategies (*4*), and (3) previous work showing state-based diversity metrics (*5*). Previous population genomic assessments have supported a hierarchical population structure in koalas consisting of five ancestral clusters (*5*) . While this is common approach to characterising population structure, it only accounts for a small proportion of the genetic variation estimated from common alleles (alleles greater than 0.05 minor allele frequency; MAF). In this study, we treat each of the 27 sample locations as spatially distinct, and independent populations to explore more nuanced changes in $N_e$ and diversity based on both relaxed and stringent MAF parameter settings (Fig. 1A; Table S1). From a demographic history standpoint, using smaller geographically distinct groups can help to reveal fine scale demographic differences, while more relaxed MAF settings can help in capturing rare alleles.

**Supplementary Information**

Historical demographic studies in koalas have used a generation time of 7 years (13), derived from the average generation time estimated for a single QLD population (6.02 years; *6*). However, generation time can vary in both time and space depending on life-history traits, environmental stochasticity, and management practices (*7*). These factors, and different collection years across populations (Table S1), suggest that VIC populations may have shorter generation times than others, particularly in the more recent past, and may explain the differences in $N_e$ trends over generational time seen in Figure 1G. Consequently, applying a uniform year-to-generation conversion across populations may yield misleading conclusions across states, and therefore generation times (not years) was employed as the most appropriate temporal metric for our reconstructions of koala demographic histories.

<u>GONE (reconstructions of recent demographic history; 0-100 generations):</u> GONE uses linkage disequilibrium over a wide range of possible recombination rates across a population to estimate the cumulative contributions of previous generations on the current generation (*8*). The result is a high resolution and accurate inference of recent demographic history (1-100 generations ago). There are several issues that need to be considered when using GONE. Primarily, ongoing gene flow between populations or substructuring within the derived population can complicate model interpretation. We minimised this issue by reducing our dataset to small spatial groups to define populations. Furthermore, GONE analyses are limited to 50,000 SNPs per chromosome for each replicate run, with the input being limited to 10 M SNPs in total and a maximum of 1 M SNPs per chromosome. If more than 50,000 SNPs are provided, then 50,000 SNPs are randomly subsampled for each chromosome for each replicate run. Despite reducing our dataset to 10 million SNPs, this dataset still managed to crash the high-performance computing system. We therefore randomly subsampled 100,000 SNPs for each chromosome by using the -shuf argument and used vcftools to generate a new vcf file containing the subsampled SNPs with the --positions argument and the subsampled list of SNPs obtained from the -shuf argument. Pseudo chromosome 4 was twice as long as the others, so we subsampled 150,000 SNPs to account for this

difference. This subsampling approach ensured the program could run effectively on the high-performance computing system while also capturing genome wide variation. Using plink, we created ped and map files for each population, and finally, we used the script_GONE.sh script provided on the github site (https://github.com/esrud/GONE). The GONE script calculates the geometric mean of $N_e$ across 40 replicate runs. Analyses were performed mostly using default input parameters. However, because of intensive recent management of koala populations the -hc parameter (highest recombination rate value) was reduced to 0.01 to lessen the effect of human induced mixing between populations on GONE outputs, as advised by the program developers. We contrasted outputs from each population to estimate variance within each state and used line plots to present the relationship between generation and mean $N_e$ using ggplot2. The geom_smooth argument was used to obtain the mean across populations along with the 95% confidence intervals.

<u>ROH (reconstructing intermediate demographic history; 0-1,000 generations):</u> We followed the analyses and outputs from Khan et al (9) to relate ROH sizes to $N_e$ through time. ROH calculations were performed using the -roh argument in bcftools, including ROHs greater than 500 kb. To estimate $N_e$ over different time frames, we used $F_{ROH}$ values derived from ROH corresponding to ancestors within several time intervals. Over time, the length distribution of ROH reflects historical changes in $N_e$. Short ROHs indicate ancient inbreeding or small $N_e$ events that occurred many generations ago and have been progressively broken down by recombination over time. In contrast, long ROHs reflect recent inbreeding or recent reductions in $N_e$, such as bottleneck events (*10, 11*). Assuming that the estimated coalescent times for ROH are unbiased, the average $F_{ROH}$ calculated from ROH with maximum coalescent times less than t generations, represents the inbreeding accumulated in the population from the sampling time back to t generations ago. We derived $N_e$ using the relationship between the mean expected individual inbreeding ($F_{ROH}$) and $N_e$ over *t* generations (9) :

$$F_{\mathrm{ROH,t}} = (1 - \frac{1}{2}\, Ne)^{t}$$

**Supplementary Information**

SMC++ (reconstructing historical demographic history; 100 to 100,000 generations): SMC++ uses a multiple sequentially Markovian coalescent model (MSMC) across unphased genomes from a population. Thus, it is more accurate than other MSMC methods, especially when estimating population sizes in the recent past (100-500 generations ago) and when phasing error is present (*12*). We used the dataset filtered by depth, quality and missing data (Table S2) for SMC++. We then used the vcf2smc script to create 11 separate input smc files for each population (one for each pseudo chromosome). The estimate function was used separately for each chromosome using a mutation rate of $1.45 \times 10^{-8}$ (*13*) and the cubic spline argument. Performing SMC++ for each chromosome allowed us to visualise the variance within populations. We then used the SMC++ plotting function for all pseudo chromosomes for each population to obtain csv files, which were imported into R to produce graphical outputs for each population using ggplot2. For each population, we used the geom_smooth argument with the -loess function across all 11 pseudo chromosomes to estimate the mean change in $N_e$ with 95% confidence intervals between 100 and 100,000 generations. In humans, MSMC methods have been shown to be accurate to around 80 generations ago (*14*). We therefore cut off visualisation at 100 generations and relied on GONE estimates for interpretations of more recent shifts in $N_e$.

Contemporary $N_e$ – Contemporary $N_e$ was estimated using the currentNe software (*15*). This software is designed to estimates $N_e$ in problematic scenarios involving variable population sizes and makes use of a small artificial neural network to estimate confidence intervals, which allows for better estimation of confidence intervals when relying on small sample sizes. Critically, this method is less biased than those used previously to re-construct the past demography (*15*). We used a default -k, which allowed the algorithm to estimate the average number of full siblings from the input data. Analyses were performed on 100,000 randomly subsampled SNPs, and we report the $N_e$ estimated and averaged across each within-chromosome calculation.

Autosomal heterozygosity and rare variants - Autosomal heterozygosity was calculated on the full dataset that was filtered only on read depth and quality and the retention of biallelic SNPs (Table S2). While these parameter settings maximise read accuracy, they do not completely control for error. However, errors are expected to be randomly distributed across all samples and are not expected to affect between-population heterozygosity signals across the genome, if anything the noise from error would depress the signal of the evolutionary pattern. To understand how rare variant diversity differed among populations, we created three dataset bins: common variants (MAF > 0.05), low frequency variants (0.05 > MAF > 0.01), and rare variants (0.01 > MAF) and explored patterns of heterozygosity within each MAF bin. These bins were created using the bcftools -view function with and combinations of -i and -e arguments to include or exclude variants at specific MAFs across the whole dataset. We used the bcftools -stat function to obtain per sample counts of heterozygotes and divided each by the combined length of the 11 pseudo chromosomes used in this study. Outputs were statistically compared using the lm function in R to identify significant models and posthoc Tukey's tests to identify significant differences between groups. These results were plotted using the boxplot function in R.

To determine if rare variants have accumulated during post-bottleneck expansion, we contrasted the ratio of autosomal heterozygosity between MAF bins. This ultimately assesses the rate of change of heterozygosity as a function of allele frequency. The theory here is that alleles with lower frequency will be lost first during a population contraction event; therefore, we would expect to find a greater loss of low and rare frequency alleles in bottlenecked populations. Conversely, the expectation is that lower frequency alleles will be gained first during a demographic expansion. We used the ratio of functional variants between MAF bins to compare the rate of change of variants per sample. If no bottleneck occurred the low/common and rare/low ratios should be the same across all populations and states. If one state has been exposed to a more severe bottleneck without expansion, we would expect that both ratios would be lower than the other groups. Given VIC has undergone rapid population expansion after a severe bottleneck we expected the rare/low ratio to be similar to other states but the low/common ratio to be much lower.

**Supplementary Information**

Inbreeding - We estimated inbreeding using three different methods. The first was a classic individual based method implemented in PLINK, which employs the method-of-moments *F* coefficient estimated using the -ibc argument. Second, we used ROHs to calculate inbreeding ($F_{ROH}$) and included any ROH greater than 500 kb. Runs of homozygosity were estimated using the bcftools -roh function with the -G30 and -r flags. Recent work shows that this provides a more accurate genomic measure of inbreeding than more traditional approaches (*16*). Lastly, we use the roh-selfing method (*17*) using the sequential random forest model to understand the theoretical selfing rates within populations (i.e., inbreeding). This software was developed to understand selfing in plants, but in purely outcrossing species such as koalas, we can directly relate theoretical selfing to inbreeding within populations. The method combines ROH, Tajima's D and *F* stats to estimate selfing rates over chromosomes. It uses random forests to identify unique characteristics of ROHs that are indicative of different sources of inbreeding. In this case, we use the term theoretical selfing rate from a genomic perspective (a different measure of inbreeding), quantifying the proportion of a genome that is related in a population.

Functional genomics - We used Variant Effect Predictor (VEP) to obtain general information about the effect of variants within and among populations (*18*). We were interested in comparing loss of function (LOF), missense (nonsynonymous variants) and regulatory region variants (5kb upstream or downstream of a gene) among populations. We focussed on these variants because they are likely to have consequential effects (positive or negative). LOF variants included splice acceptor variants, splice donor variants, stop gained variants, and splice region variants. Frameshift variants were not included as insertions and deletions were filtered out prior to analysis. Because the number of variants changed between populations, and the reference genome represents a northern population, we standardised the number of consequential variants using the $R_{X/Y}$ statistic previously developed (*19*) and used in several recent studies (*9*, *20*, *21*). We follow the methods from (*21*) because, like their work, we were unable to accurately infer the ancestral allele in koalas. Briefly, the $R_{X/Y}$ method compares groups to test if there is a higher rate of functional variants (LOF, missense, and regulatory region) compared to the number of intergenic sites. At each site i, the non-reference allele frequency in population X and Y, was determined using bcftools. Then $L_{X,Y}$ was estimated for each functional category using the following formulas, where the numerator is the consequential variant (C) and the denominator is the intergenic variant (I):

$$L_{X,Y}(C) = \frac{\sum_{i \in C} f_i^X (1 - f_i^Y)}{\sum_{j \in I} f_j^X (1 - f_j^Y)}$$

$$L_{Y,X}(C) = \frac{\sum_{i \in C} f_i^Y (1 - f_i^X)}{\sum_{j \in I} f_j^Y (1 - f_j^X)}$$

The sum of the intergenic sites (I) mitigates the population-specific reference mapping biases. And then $R_{X/Y}$ is estimated as the ratio of these two consequential ratios:

$$R_{X/Y} = \frac{L_{X,Y}(C)}{L_{Y,X}(C)}$$

Finally, we used 100 block jackknifes on the set of sites in C to determine the 95% confidence intervals for each $R_{X/Y}$ estimate. The confidence intervals were very small because the blocked data sets were still very large (not shown in the figure but given in Table S3). Results are given in the form of barplots which were generated using ggplot2. $R_{X/Y}$ is interpreted as an excess of the consequential variant type for the population (X) in the numerator if the number is greater than 1 or the population in the denominator (Y) if the number is less than 1.

Recent works discuss the impact of LOF variants and assume that they are deleterious and termed 'genetic load'. However, this is an oversimplification of LOF variants and an unreliable indicator of genetic load. LOF variants are often assumed to have severe consequences, being deleterious and strongly selected against (*22*). While purifying selection indeed biases LOF variants toward low allele frequencies (*23–25*), several studies indicate that many LOF variants are benign (*26*), support protein homeostasis (*27*), can be beneficial (*22, 28*) and contribute to adaptive processes (*29*). Additionally, bacterial systems demonstrate that LOF in cryptic operons can be reversed under environmental shifts (*30, 31*). Factors such as genetic redundancy, paralogs, gene essentiality, pseudogenes, and LOF variants present in reference genomes can also inflate estimates of deleterious LOF variants, complicating their interpretation. Thus, linking LOF variants directly to genetic load in population genetic studies remains tenuous. Estimating true genetic load would require temporal studies on population dynamics, fitness, and behaviour. Even so, the differential accumulation of these variants is an effective way to highlight the consequential effects of genetic drift and the recent accumulation of functional variants, without interpreting them as beneficial or deleterious. We also present the distribution of missense and regulatory variants between populations because they are known to be important in adaptation among species across kingdoms (*32–34*). It is important to acknowledge that these variants could be beneficial, neutral, or deleterious and their impacts could be environmentally dependent (deleterious in one environment while beneficial in another; (*35*).

To ascertain the differences in genetic diversity between state meta-populations, we also explored functional variant differences in state specific (private) alleles. Private alleles can be broadly implicated in adaptive processes (*36*), can substantiate the effect of genetic drift (*14*), and help to illustrate broad genetic differences between states (*37*). We used custom R code on a combination of the VEP outputs with state specific allele frequencies calculated from bcftools. To standardise this output, we divided the total number of private variants in the specific bin by the total number of samples. We also explored the interaction of purging across MAF bins and functional groups. This was achieved by contrasting the change in functional alleles per sample across MAF bins to determine if they change across states. Results are graphically depicted in the form of a line plot generated using ggplot2.

### References

1. C. J. Hogg, L. Silver, E. A. McLennan, K. Belov, Koala Genome Survey: An Open Data Resource to Improve Conservation Planning. *Genes-basel* **14**, 546 (2023).

2. R. O. Betschart, A. Thiéry, D. Aguilera-Garcia, M. Zoche, H. Moch, R. Twerenbold, T. Zeller, S. Blankenberg, A. Ziegler, Comparison of calling pipelines for whole genome sequencing: an empirical study demonstrating the importance of mapping and alignment. *Sci. Rep.* **12**, 21502 (2022).

3. S. Zhao, O. Agafonov, A. Azab, T. Stokowy, E. Hovig, Accuracy and efficiency of germline variant calling pipelines for human genome data. *Sci. Rep.* **10**, 20222 (2020).

4. P. Menkhorst, Too close for comfort. *In* Contentious issues in human-wildlife encounters, edited by D. Lunney, A. Munn and W. Meikle. Royal Zoological Society of New South Wales, Mosman, NSW, Australia. 73–92 (2008).

5. E. A. McLennan, T. G. L. Kovacs, L. W. Silver, Z. Chen, F. R. Jaya, S. Y. W. Ho, K. Belov, C. J. Hogg, Genomics identifies koala populations at risk across eastern Australia. *Ecol. Appl.*, e3062 (2024).

6. S. S. Phillips, Population Trends and the Koala Conservation Debate. *Conserv. Biol.* **14**, 650–659 (2000).

7. Y. G. Araya-Ajoy, G. H. Bolstad, J. Brommer, V. Careau, N. J. Dingemanse, J. Wright, Demographic measures of an individual's "pace of life": fecundity rate, lifespan, generation time, or a composite variable? *Behav. Ecol. Sociobiol.* **72**, 75 (2018).

8. E. Santiago, I. Novo, A. F. Pardiñas, M. Saura, J. Wang, A. Caballero, Recent Demographic History Inferred by High-Resolution Analysis of Linkage Disequilibrium. *Mol. Biol. Evol.* **37**, 3642–3653 (2020).

9. A. Khan, K. Patel, H. Shukla, A. Viswanathan, T. van der Valk, U. Borthakur, P. Nigam, A. Zachariah, Y. V. Jhala, M. Kardos, U. Ramakrishnan, Genomic evidence for inbreeding depression and purging of deleterious genetic variation in Indian tigers. *Proc. Natl. Acad. Sci.* **118**, e2023018118 (2021).

10. A. D. Foote, R. Hooper, A. Alexander, R. W. Baird, C. S. Baker, L. Ballance, J. Barlow, A. Brownlow, T. Collins, R. Constantine, L. D. Rosa, N. J. Davison, J. W. Durban, R. Esteban, L. Excoffier, S. L. F. Martin, K. A. Forney, T. Gerrodette, M. T. P. Gilbert, C. Guinet, M. B. Hanson, S. Li, M. D. Martin, K. M. Robertson, F. I. P. Samarra, R. de Stephanis, S. B. Tavares, P. Tixier, J. A. Totterdell, P. Wade, J. B. W. Wolf, G. Fan, Y. Zhang, P. A. Morin, Runs of homozygosity in killer whale genomes provide a global record of demographic histories. *Mol. Ecol.* **30**, 6162–6177 (2021).

11. A. M. Hewett, M. A. Stoffel, L. Peters, S. E. Johnston, J. M. Pemberton, Selection, recombination and population history effects on runs of homozygosity (ROH) in wild red deer (Cervus elaphus). *Heredity* **130**, 242–250 (2023).

12. J. Terhorst, J. A. Kamm, Y. S. Song, Robust and scalable inference of population history from hundreds of unphased whole genomes. *Nat. Genet.* **49**, 303–309 (2017).

13. R. N. Johnson, D. O'Meally, Z. Chen, G. J. Etherington, S. Y. W. Ho, W. J. Nash, C. E. Grueber, Y. Cheng, C. M. Whittington, S. Dennison, E. Peel, W. Haerty, R. J. O'Neill, D. Colgan, T. L. Russell, D. E. Alquezar-Planas, V. Attenbrow, J. G. Bragg, P. A. Brandies, A.

Y.-Y. Chong, J. E. Deakin, F. D. Palma, Z. Duda, M. D. B. Eldridge, K. M. Ewart, C. J. Hogg, G. J. Frankham, A. Georges, A. K. Gillett, M. Govendir, A. D. Greenwood, T. Hayakawa, K. M. Helgen, M. Hobbs, C. E. Holleley, T. N. Heider, E. A. Jones, A. King, D. Madden, J. A. M. Graves, K. M. Morris, L. E. Neaves, H. R. Patel, A. Polkinghorne, M. B. Renfree, C. Robin, R. Salinas, K. Tsangaras, P. D. Waters, S. A. Waters, B. Wright, M. R. Wilkins, P. Timms, K. Belov, Adaptation and conservation insights from the koala genome. *Nat. Genet.* **50**, 1102–1111 (2018).

14. N. Mather, S. M. Traves, S. Y. W. Ho, A practical introduction to sequentially Markovian coalescent methods for estimating demographic history from genomic data. *Ecol. Evol.* **10**, 579–589 (2020).

15. E. Santiago, A. Caballero, C. Köpke, I. Novo, Estimation of the contemporary effective population size from SNP data while accounting for mating structure. *Mol. Ecol. Resour.* **24**, e13890 (2024).

16. K. T. Mekonnen, D.-H. Lee, Y.-G. Cho, A.-Y. Son, K.-S. Seo, Genomic and Conventional Inbreeding Coefficient Estimation Using Different Estimator Models in Korean Duroc, Landrace, and Yorkshire Breeds Using 70K Porcine SNP BeadChip. *Animals* **14**, 2621 (2024).

17. L. Zeitler, K. J. Gilbert, Using Runs of Homozygosity and Machine Learning to Disentangle Sources of Inbreeding and Infer Self-Fertilization Rates. *Genome Biol. Evol.* **16**, evae139 (2024).

18. W. McLaren, L. Gil, S. E. Hunt, H. S. Riat, G. R. S. Ritchie, A. Thormann, P. Flicek, F. Cunningham, The Ensembl Variant Effect Predictor. *Genome Biol.* **17**, 122 (2016).

19. R. Do, D. Balick, H. Li, I. Adzhubei, S. Sunyaev, D. Reich, No evidence that selection has been less effective at removing deleterious mutations in Europeans than in Africans. *Nat. Genet.* **47**, 126–131 (2015).

20. M. Kardos, Y. Zhang, K. M. Parsons, Y. A, H. Kang, X. Xu, X. Liu, C. O. Matkin, P. Zhang, E. J. Ward, M. B. Hanson, C. Emmons, M. J. Ford, G. Fan, S. Li, Inbreeding depression explains killer whale population dynamics. *Nat. Ecol. Evol.* **7**, 675–686 (2023).

21. Y. Xue, J. Prado-Martinez, P. H. Sudmant, V. Narasimhan, Q. Ayub, M. Szpak, P. Frandsen, Y. Chen, B. Yngvadottir, D. N. Cooper, M. de Manuel, J. Hernandez-Rodriguez, I. Lobon, H. R. Siegismund, L. Pagani, M. A. Quail, C. Hvilsom, A. Mudakikwa, E. E. Eichler, M. R. Cranfield, T. Marques-Bonet, C. Tyler-Smith, A. Scally, Mountain gorilla genomes reveal the impact of long-term population decline and inbreeding. *Science* **348**, 242–245 (2015).

22. N. Whiffin, K. J. Karczewski, X. Zhang, S. Chothani, M. J. Smith, D. G. Evans, A. M. Roberts, N. M. Quaife, S. Schafer, O. Rackham, J. Alföldi, A. H. O'Donnell-Luria, L. C. Francioli, I. M. Armean, E. Banks, L. Bergelson, K. Cibulskis, R. L. Collins, K. M. Connolly, M. Covarrubias, B. Cummings, M. J. Daly, S. Donnelly, Y. Farjoun, S. Ferriera, S. Gabriel, L. D. Gauthier, J. Gentry, N. Gupta, T. Jeandet, D. Kaplan, K. M. Laricchia, C. Llanwarne, E. V. Minikel, R. Munshi, B. M. Neale, S. Novod, N. Petrillo, T. Poterba, D. Roazen, V. Ruano-Rubio, A. Saltzman, K. E. Samocha, M. Schleicher, C. Seed, M. Solomonson, J. Soto, G.

Tiao, K. Tibbetts, C. Tolonen, et al., Characterising the loss-of-function impact of 5' untranslated region variants in 15,708 individuals. *Nat. Commun.* **11**, 2523 (2020).

23. D. G. MacArthur, S. Balasubramanian, et al., A Systematic Survey of Loss-of-Function Variants in Human Protein-Coding Genes. *Science* **335**, 823–828 (2012).

24. R. M. Durbin, D. Altshuler, R. M. Durbin, et al., A map of human genome variation from population-scale sequencing. *Nature* **467**, 1061–1073 (2010).

25. Y.-C. Xu, X.-M. Niu, X.-X. Li, W. He, J.-F. Chen, Y.-P. Zou, Q. Wu, Y. E. Zhang, W. Busch, Y.-L. Guo, Adaptation and phenotypic civersification in *Arabidopsis* through loss-of-function mutations in protein-coding genes. *Plant Cell* **31**, 1012–1025 (2019).

26. D. G. MacArthur, C. Tyler-Smith, Loss-of-function variants in the genomes of healthy humans. *Hum. Mol. Genet.* **19**, R125–R130 (2010).

27. C. L. Shovlin, M. A. Aldred, When "loss-of-function" means proteostasis burden: Thinking again about coding DNA variants. *Am. J. Hum. Genet.* **112**, 3–10 (2025).

28. J. Serrano, S. Kondo, G. M. Link, I. S. Brown, R. E. Pratley, K. K. Baskin, B. H. Goodpaster, P. M. Coen, G. A. Kyriazis, A partial loss-of-function variant (Ile191Val) of the TAS1R2 glucose receptor is associated with enhanced responses to exercise training in older adults with obesity: A translational study. *Metabolism* **162**, 156045 (2025).

29. Y.-C. Xu, Y.-L. Guo, Less Is More, Natural Loss-of-Function Mutation Is a Strategy for Adaptation. *Plant Commun.* **1**, 100103 (2020).

30. B. G. Hall, S. Yokoyama, D. H. Calhoun, Role of cryptic genes in microbial evolution. *Mol. Biol. Evol.* **1**, 109–124 (1983).

31. B. G. Hall, L. Xu, Nucleotide sequence, function, activation, and evolution of the cryptic asc operon of Escherichia coli K12. *Mol. Biol. Evol.* **9**, 688–706 (1992).

32. F. W. Albert, L. Kruglyak, The role of regulatory variation in complex traits and disease. *Nat Rev Genet* **16**, 197–212 (2015).

33. C. W. Ahrens, K. Murray, R. A. Mazanec, S. Ferguson, A. Jones, D. T. Tissue, M. Byrne, J. O. Borevitz, P. D. Rymer, Genomic determinants, architecture, and constraints in drought-related traits in Corymbia calophylla. *BMC Genom.* **25** (2024).

34. T. Latrille, N. Rodrigue, N. Lartillot, Genes and sites under adaptation at the phylogenetic scale also exhibit adaptation at the population-genetic scale. *Proc. Natl. Acad. Sci.* **120**, e2214977120 (2023).

35. J. T. ANDERSON, C. LEE, C. A. RUSHWORTH, R. I. COLAUTTI, T. MITCHELL-OLDS, Genetic trade-offs and conditional neutrality contribute to local adaptation. *Mol Ecol* **22**, 699–708 (2013).

36. A. E. Sjöstrand, P. Sjödin, M. Jakobsson, Private haplotypes can reveal local adaptation. *Bmc Genet* **15**, 61–61 (2014).

37. T. Maruki, Z. Ye, M. Lynch, Evolutionary Genomics of a Subdivided Species. *Mol. Biol. Evol.* **39**, msac152 (2022).

**Supplementary Information**

### Supplementary Tables

**Table S1**. Koala population information and summary statistics for each of the 27 populations. $N_e$ is the contemporary estimate of $N_e$ using currentNe from LD between chromosomes. Individual *F* from plink using the moments of methods method.

| Population | State | n | TajD (SE) | $F_{ROH}$ > 500kb | *F* | $N_e$ | *90%CI (LL-UL)* |
|---|---|---|---|---|---|---|---|
| BLMT | NSW | 15 | 0.85 (0.005) | 0.28 | 0.23 | 10.00 | 7.80-12.81 |
| BURR | QLD | 16 | 0.56 (0.004) | 0.15 | -0.02 | 243.51 | 107.30-552.63 |
| CAMT | NSW | 46 | 1.55 (0.007) | 0.37 | 0.37 | 21.92 | 20.33-23.64 |
| CLAV | NSW | 5 | 1.56 (0.004) | 0.15 | 0.02 | 77.20 | 32.84-181.48 |
| COTW | VIC | 5 | 0.65 (0.008) | 0.58 | 0.65 | 855.60 | 115.44-6341.58 |
| EGIP | VIC | 11 | 1.22 (0.009) | 0.63 | 0.69 | 89.60 | 56.23-142.80 |
| FRAS | QLD | 10 | 0.64 (0.005) | 0.31 | 0.15 | 14.80 | 11.37-19.27 |
| FREI | VIC | 2 | 0.12 (0.007) | 0.57 | 0.63 | NA | NA |
| GILB | QLD | 3 | 0.21 (0.005) | 0.16 | -0.03 | 91.6 | 23.03-364.29 |
| GOLD | QLD | 17 | 0.60 (0.004) | 0.22 | 0.09 | 138.52 | 92.95-206.45 |
| KYOG | NSW | 25 | 0.61 (0.004) | 0.16 | 0.05 | 101.99 | 79.90-130.19 |
| MALL | VIC | 7 | 0.75 (0.009) | 0.62 | 0.68 | 55.20 | 31.17-97.76 |
| MONA | NSW | 21 | 1.42 (0.007) | 0.27 | 0.22 | 98.80 | 73.93-132.04 |
| MORB | QLD | 16 | 0.89 (0.005) | 0.19 | 0.46 | 244.95 | 148.95-402.85 |
| MURR | VIC | 12 | 1.17 (0.009) | 0.59 | 0.65 | 82.80 | 54.39-126.05 |
| NARR | NSW | 17 | 1.27 (0.005) | 0.28 | 0.27 | 68.40 | 51.78-90.36 |
| NRIV | NSW | 48 | 1.15 (0.005) | 0.26 | 0.18 | 49.85 | 44.92-55.31 |
| PILL | NSW | 38 | 0.54 (0.005) | 0.30 | 0.23 | 19.20 | 17.61-20.93 |
| PMAC | NSW | 17 | 0.72 (0.005) | 0.18 | 0.73 | 60.50 | 45.40-80.61 |
| PTS | NSW | 10 | 0.70 (0.007) | 0.36 | 0.28 | 26.93 | 19.57-37.06 |
| QLDM | QLD | 2 | 0.21 (0.005) | 0.20 | 0.02 | 32.32 | 8.70-120.02 |
| REDL | QLD | 5 | 0.49 (0.005) | 0.34 | 0.22 | 4.94 | 5.01-6.64 |
| SGIP | VIC | 24 | 1.00 (0.01) | 0.51 | 0.56 | 69.60 | 47.29-102.42 |
| SHBS | VIC | 7 | 0.80 (0.008) | 0.54 | 0.60 | 226.17 | 83.91-609.51 |
| SUNC | QLD | 15 | 0.58 (0.004) | 0.19 | 0.04 | 122.03 | 81.09-183.64 |
| TOOW | QLD | 20 | 0.64 (0.004) | 0.20 | 0.06 | 249.35 | 161.22-385.65 |
| WVIC | VIC | 4 | 0.50 (0.008) | 0.55 | 0.62 | 98.00 | 31.87-301.39 |

#### Supplementary Information

**Table S2.** Details of the data filtering applied to generate datasets for each analysis undertaken in the current study

| Analysis | Measure | PHRED | Read depth | Missing data | Minor allele frequency | Minor allele count | Subsample |
|---|---|---|---|---|---|---|---|
| ROH | Inbreeding, Ne, intermediate demography, theoretical selfing | 30 | >10; <100 | 0.05 | 0.01 | | |
| Autosomal heterozygosity | diversity | 30 | >10; <100 | 0.05 | | | |
| Rare alleles | general | 30 | >10; <100 | 0.05 | | | |
| SMC++ | Historical demography | 30 | >10; <100 | 0.05 | 0.01 | | |
| GONE | Recent demography | 30 | >10; <100 | 0.05 | 0.05 | | 100-150k SNPs per chromosome |
| currentNe | Current demography | 30 | >10; <100 | 0.05 | 0.01 | | 100k SNPs per chromosome |
| Functional | VEP, $R_{XY}$, private alleles, @ population level | 30 | >10; <100 | 0.05 | | 1 | |

#### Supplementary Information

**Table S3.** Estimated statistics for $R_{X/Y}$ analysis. MAF group = minor allele frequency group. Function = functional group (LOF = loss of function; missense = nonsynonymous; regulatory = variants within 5kb of a gene). 95% confidence interval was calculated from 100 blocks similar to the method performed in (*21*) along with significance values.

| MAF group | function | numerator | denominator | $R_{X/Y}$ | 95% CI | Z-score | p-value |
|---|---|---|---|---|---|---|---|
| common | LOF | NSW | QLD | 1.037 | 0.0004 | 20.7 | < 0.001 |
| common | LOF | NSW | VIC | 0.929 | 0.0005 | -25.7 | < 0.001 |
| common | LOF | QLD | VIC | 0.912 | 0.0006 | -31.2 | < 0.001 |
| common | missense | NSW | QLD | 1.030 | 0.0001 | 38.5 | < 0.001 |
| common | missense | NSW | VIC | 0.973 | 0.0002 | -23.3 | < 0.001 |
| common | missense | QLD | VIC | 0.953 | 0.0002 | -38.1 | < 0.001 |
| common | regulatory | NSW | QLD | 1.019 | < 0.0001 | 155.1 | < 0.001 |
| common | regulatory | NSW | VIC | 0.965 | < 0.0001 | -180.3 | < 0.001 |
| common | regulatory | QLD | VIC | 0.957 | < 0.0001 | -198.8 | < 0.001 |
| low | LOF | NSW | QLD | 1.028 | 0.0006 | 9.5 | < 0.001 |
| low | LOF | NSW | VIC | 0.836 | 0.001 | -26.9 | < 0.001 |
| low | LOF | QLD | VIC | 0.822 | 0.001 | -26.6 | < 0.001 |
| low | missense | NSW | QLD | 1.063 | 0.0003 | 40.1 | < 0.001 |
| low | missense | NSW | VIC | 0.801 | 0.0004 | -80.1 | < 0.001 |
| low | missense | QLD | VIC | 0.774 | 0.0004 | -107.7 | < 0.001 |
| low | regulatory | NSW | QLD | 0.997 | < 0.0001 | -14.5 | < 0.001 |
| low | regulatory | NSW | VIC | 0.887 | 0.0001 | -185.6 | < 0.001 |
| low | regulatory | QLD | VIC | 0.801 | < 0.0001 | -210.0 | < 0.001 |
| rare | LOF | NSW | QLD | 0.999 | 0.0007 | -0.37 | 0.354 |
| rare | LOF | NSW | VIC | 0.915 | 0.0017 | -9.8 | < 0.001 |
| rare | LOF | QLD | VIC | 0.912 | 0.0015 | -10.9 | < 0.001 |
| rare | missense | NSW | QLD | 0.979 | 0.0002 | -18.4 | < 0.001 |
| rare | missense | NSW | VIC | 0.770 | 0.0004 | -96.1 | < 0.001 |
| rare | missense | QLD | VIC | 0.801 | 0.0004 | -79.6 | < 0.001 |
| rare | regulatory | NSW | QLD | 0.975 | < 0.0001 | -81.4 | < 0.001 |
| rare | regulatory | NSW | VIC | 0.889 | 0.0001 | -162.6 | < 0.001 |
| rare | regulatory | QLD | VIC | 0.919 | 0.0001 | -119.5 | < 0.001 |

### Supplementary Figures

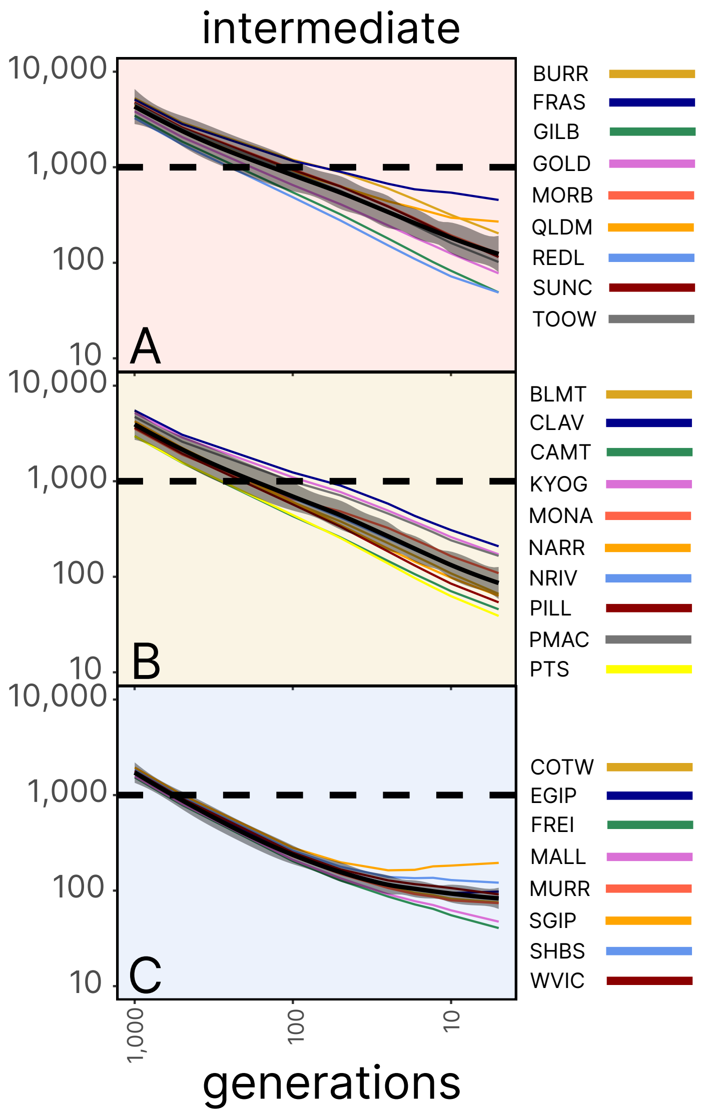


**Fig S1.** Reconstruction of effective population size ($N_e$) over intermediate time scales based on analysis of runs of homozygosity (ROH).

**Supplementary Information**

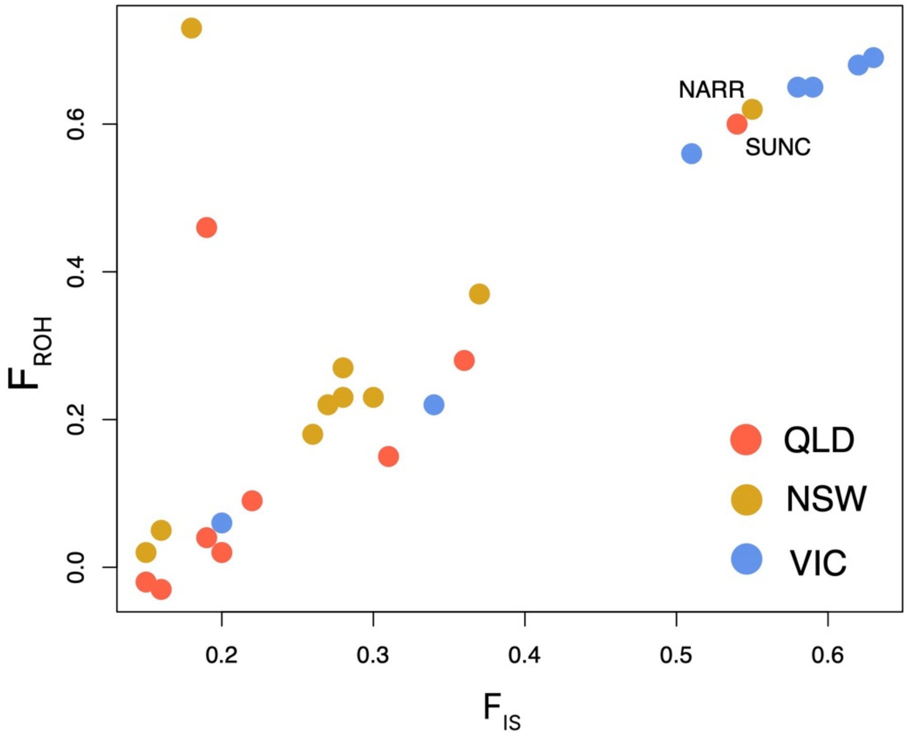


**Fig S2.** Scatter plot demonstrating the relationship between the two individual measures of inbreeding.

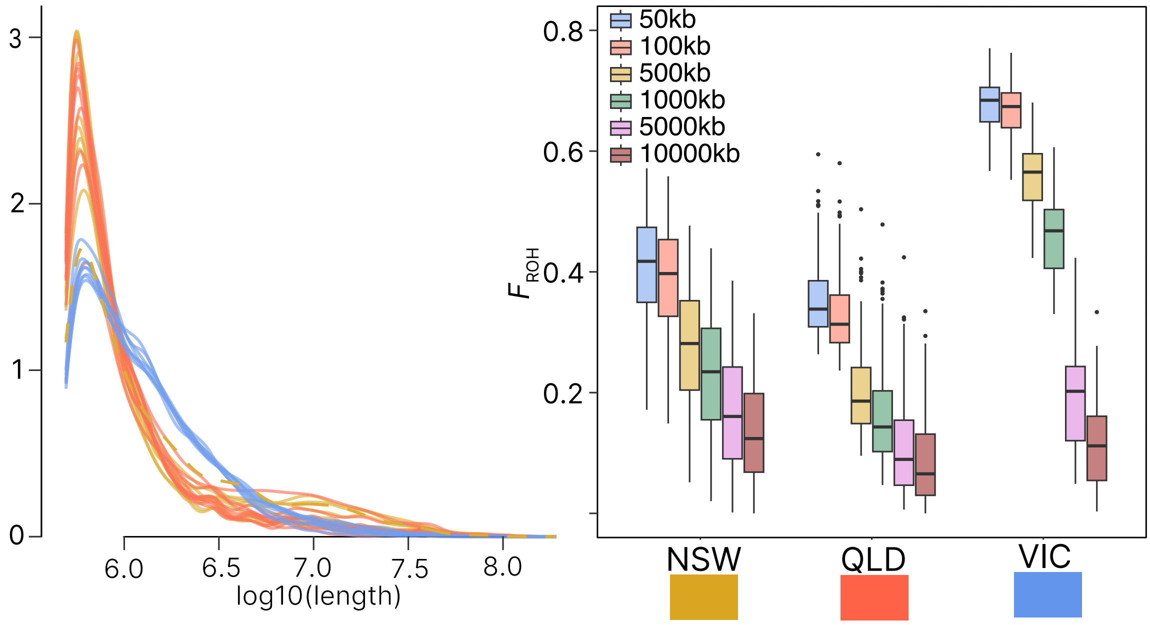


**Fig S3.** Changes in runs of homozygosity (ROH) and inbreeding based on the proportion of the genome as a ROH ($F_{ROH}$). (Left) Density plot showing differences of ROH among individuals colored by state and (right) is the differences between cumulative ROH bins for each state.

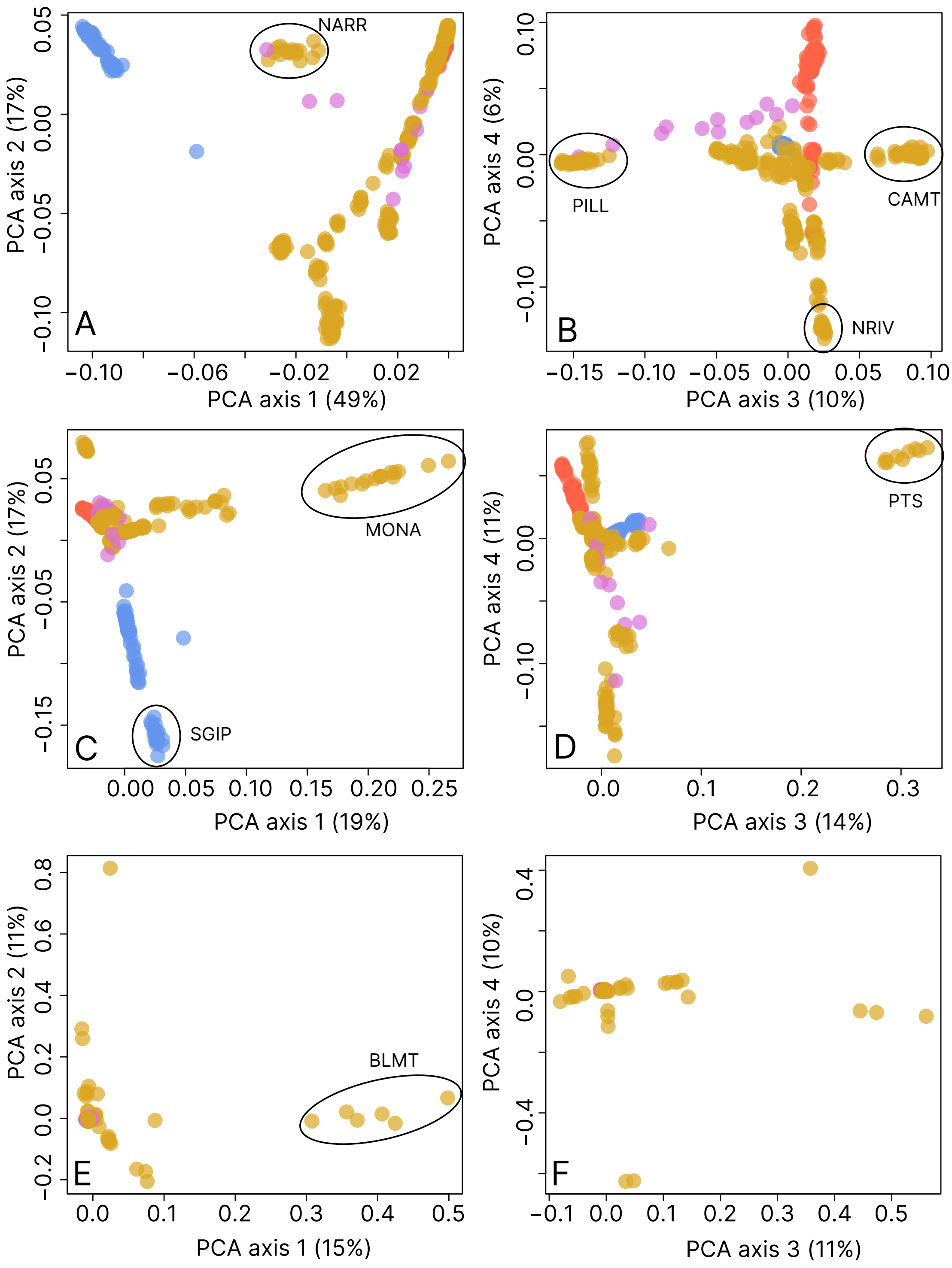


**Fig S4.** Principal coordinate analyses for each MAF bin. (A,B) represent the four axes associated with the common MAF bin. (C,D) represent the four axes for the low MAF bin. (E,F) represent the four axes for the rare MAF bin.

**Supplementary Information**

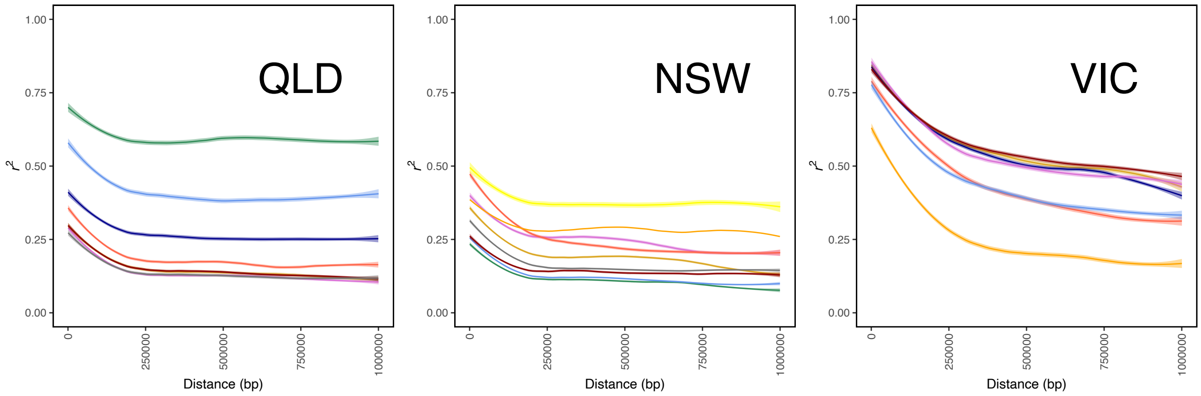


**Fig S5.** Estimates of linkage disequilibrium decay for each population.
